## Supplementary material for "Blocking SHP2 benefits FGFR2 inhibitor and overcomes its resistance in FGFR2-amplified gastric cancer": Supplrmentary data

| | No.of patients | With FGFR2 amplification n=10(%) | Without FGFR2 amplification n=151(%) | P[a] |
|---|---|---|---|---|
| Age(years) | | | | 0.9838 |
| ＜60 | 81 | 5(50%) | 76(50.3%) | |
| ≥60 | 80 | 5(50%) | 75(49.7%) | |
| Gender | | | | 0.9474 |
| Male | 95 | 6(60%) | 89(58.9%) | |
| Female | 66 | 4(40%) | 62(41.1%) | |
| AJCC | | | | 0.4026 |
| I-II | 9 | 0(0%) | 9(7.3%) | |
| III-IV | 124 | 9(100%) | 115(92.7%) | |
| T stage | | | | 0.8825 |
| 1-3 | 52 | 4(57.1%) | 48(60%) | |
| 4a-4b | 35 | 3(42.9%) | 32(40%) | |
| N stage | | | | 0.4404 |
| 0-2 | 28 | 1(20%) | 27(35.1%) | |
| 3a-3b | 54 | 4(80%) | 5O(64.9%) | |
| PD-L1 | | | | 0.4672 |
| positive | 40 | 2(33.3%) | 38(48.7%) | |
| negative | 44 | 4(66.7%) | 40(51.3%) | |
| MSI status | | | | 0.4537 |
| MSI-H | 8 | 0(0%) | 8(5.3%) | |
| MSS | 152 | 10(100%) | 142(94.7%) | |

AJCC American Joint Committee on Cancer

[a]Chi-square test was used in statical analyses.

**Table S1.** Clinical characteristics of GC patients in Nanjing Drum Tower Hospital cohort with and without FGFR2 amplification.

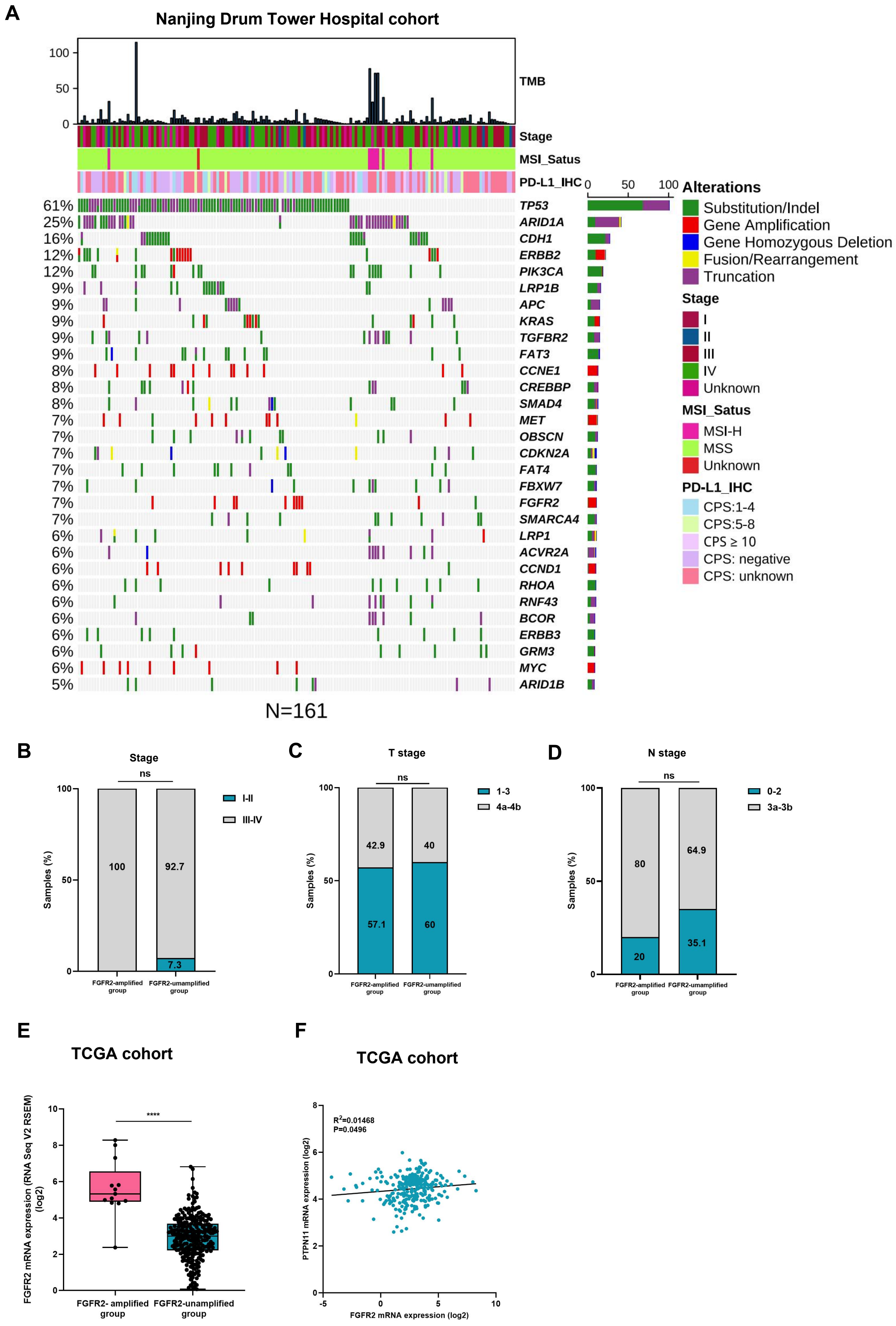


**Figure S1.**
(A) Overview of genomic alterations in GC patient samples collected in Nanjing Drum Tower Hospital (n=161). The patient samples are shown on the x-axis. Information of TMB, stage, MSI-status, CPS score and significantly altered genes are shown on the y-axis, with frequency of each alteration annotated on the right of the waterfall plot. Proportions of (B) different AJCC stages, (C,D) TNM stages among FGFR2-amplified group and FGFR2-unamplified group from Nanjing Drum Tower hospital cohort. (E) FGFR2 mRNA expression levels were analyzed between FGFR2-amplified group (n=13) and FGFR2-unamplified group (n=250) from TCGA-STAD cohort. (F) Correlation between PTPN11 and FGFR2 mRNA expressions among samples from TCGA-STAD hohort. Data are shown as Mean±SEM. ns, not significant, ****p < 0.0001. P values are detemined by Pearson's Chi-square test, linear regression t test or Fisher's exact test.

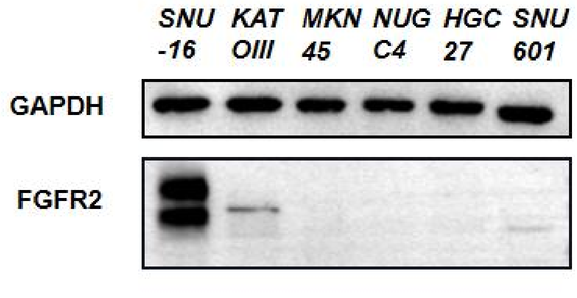


**Figure S2.** Expression levels of total-FGFR2 in diferent human GC cell lines were deteced by western blotting.

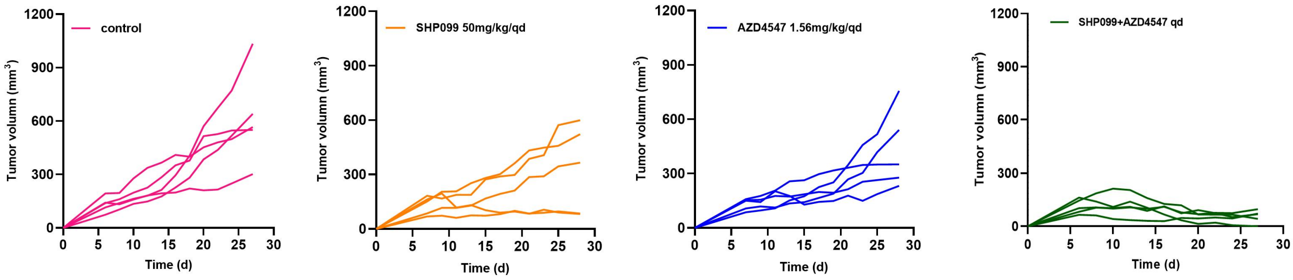


**Figure S3.** Tumor volume of individual mice in control, SHP099, AZD4547, SHP099+AZD4547 group.

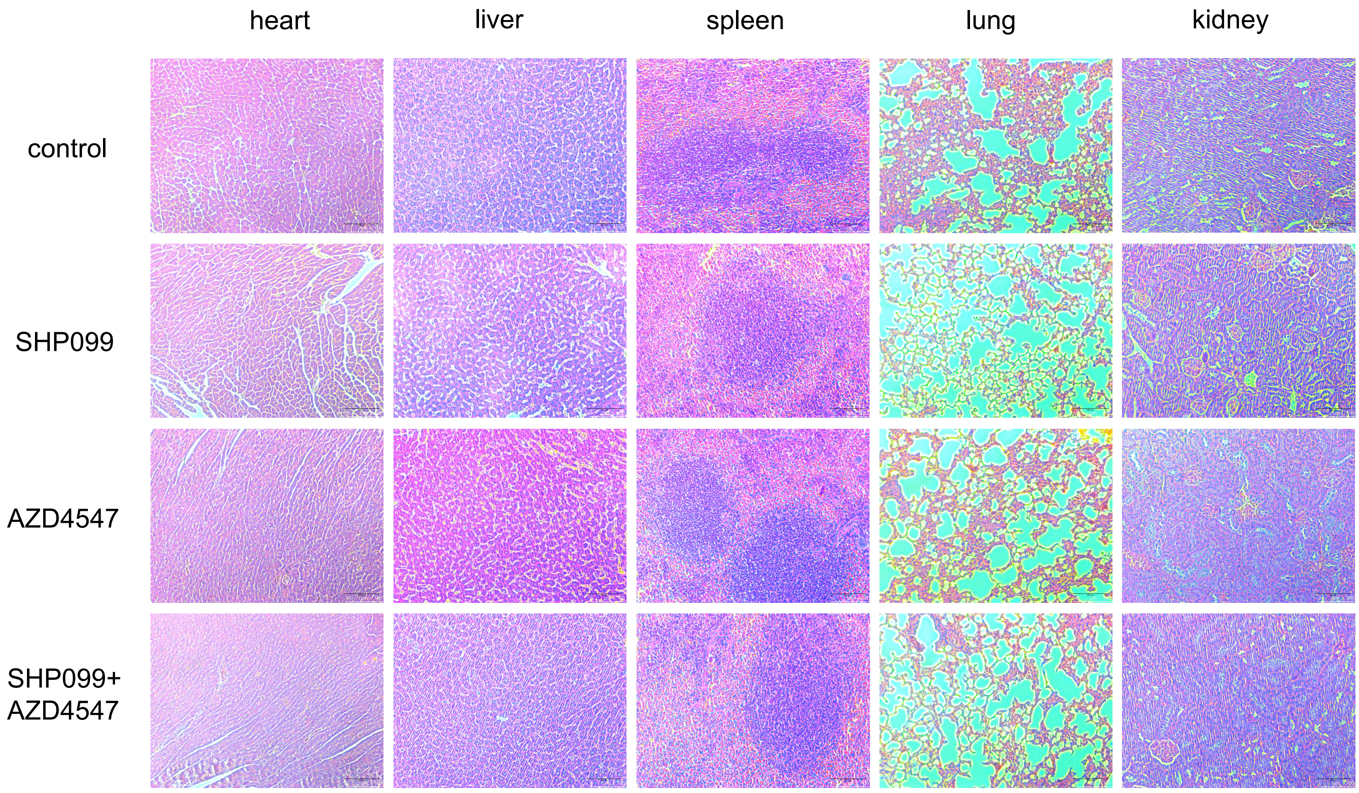


**Figure S4.** SHP2 inhibition combined with FGFR2 inhibition is safe in SNU-16 xenograft nude mice model. Safety evaluations of different administrations of mice organs including heart, liver, spleen, lung, kidney are shown by hematoxylin and eosin (H&E) staining (Scale bars, 100 μm).

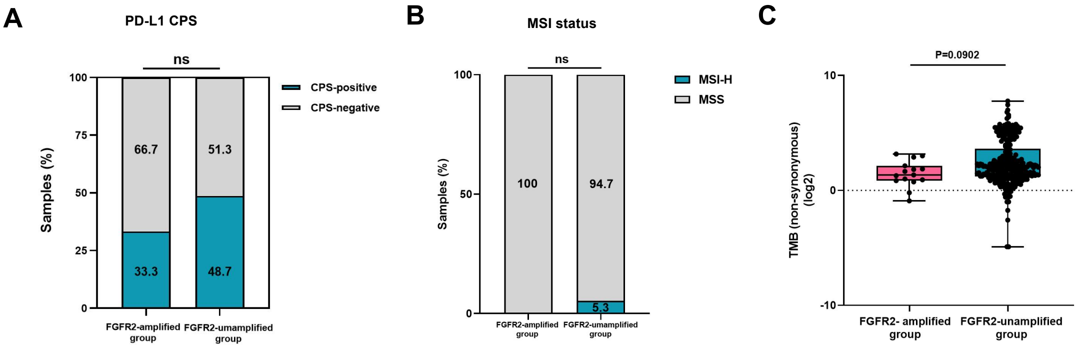


**Figure S5.** (A) PD-L1 CPS outcomes and (B) MSI status among FGFR2-amplified group and FGFR2-unamplified group from Nanjing Drum Tower hospital cohort. (C) TMB levels were analyzed between FGFR2-amplified group (n=15) and FGFR2-unamplified group (n=274) from TCGA-STAD cohort. Data are shown as Mean±SEM. ns, not significant, P values are detemined by Wilcoxon Test, Pearson's Chi-square test or Fisher's exact test. CPS≥1 was defined as CPS-positive and CPS < 1 was defind as CPS-negative.

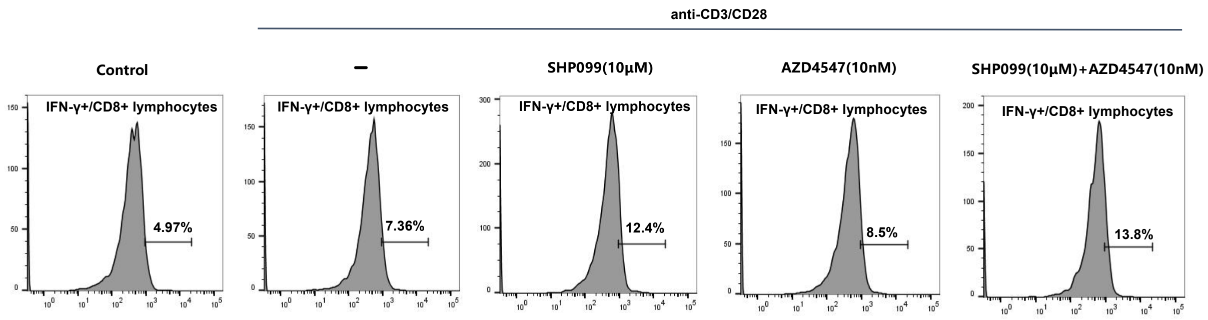


**Figure S6.** Human peripheral blood mononuclear cells (PBMCs) were incubated with different administrations in the presence of 0.25 μg/ml human anti-CD3 and 1 μg/ml human anti-CD28. Proportions of IFN-γ/CD8⁺ cells were detected by flow cytometry assay after 24-hour of drugs incubation.

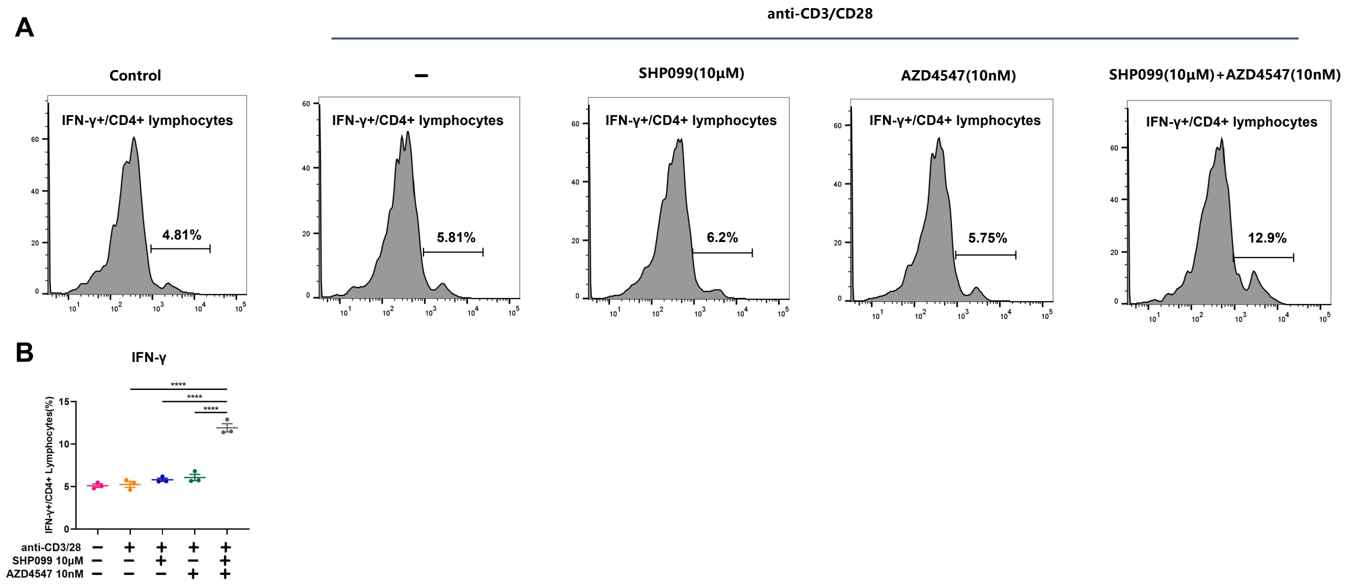


**Figure S7.** Human peripheral blood mononuclear cells (PBMCs) were incubated with different administrations in the presence of 0.25 μg/ml human anti-CD3 and 1 μg/ml human anti-CD28 for 24 hours. (A, B) Proportions of IFN-γ/CD4+ cells were detected by flow cytometry assay. Data are shown as Mean±SEM. ****p < 0.0001. P values are detemined by ordinary one-way ANOVA.
